# Psilocin acutely disrupts sleep and affects local but not global sleep homeostasis in laboratory mice

**DOI:** 10.1101/2021.02.16.431276

**Authors:** Christopher W. Thomas, Cristina Blanco-Duque, Benjamin Bréant, Guy M. Goodwin, Trevor Sharp, David M. Bannerman, Vladyslav V. Vyazovskiy

**Author notes:** Christopher Thomas is currently employed by COMPASS Pathways, plc, however, the experimental work described within was completed prior to this. The authors declare no other competing interests, financial or otherwise.

## Abstract

Serotonergic psychedelic drugs, such as psilocin (4-hydroxy-N,N-dimethyltryptamine), profoundly alter the quality of consciousness through mechanisms which are incompletely understood. Growing evidence suggests that a single psychedelic experience can positively impact long-term psychological well-being, with relevance for the treatment of psychiatric disorders, including depression. A prominent factor associated with psychiatric disorders is disturbed sleep, and the sleep-wake cycle is implicated in the regulation of neuronal firing and activity homeostasis. It remains unknown to what extent psychedelic agents directly affect sleep, in terms of both acute arousal and homeostatic sleep regulation. Here, chronic *in vivo* electrophysiological recordings were obtained in mice to track sleep-wake architecture and cortical activity after psilocin injection. Administration of psilocin led to delayed REM sleep onset and reduced NREM sleep maintenance for up to approximately 3 hours after dosing, and the acute EEG response was associated primarily with an enhanced oscillation around 4 Hz. No long-term changes in sleep-wake quantity were found. When combined with sleep deprivation, psilocin did not alter the dynamics of homeostatic sleep rebound during the subsequent recovery period, as reflected in both sleep amount and EEG slow wave activity. However, psilocin decreased the recovery rate of sleep slow wave activity following sleep deprivation in the local field potentials of electrodes targeting medial prefrontal and surrounding cortex. It is concluded that psilocin affects both global vigilance state control and local sleep homeostasis, an effect which may be relevant for its antidepressant efficacy.

## Introduction

Psilocybin is a classical serotonergic psychedelic; a unique class of drugs capable of inducing profound alterations of perception, cognition, and behaviour, commonly characterised as the psychedelic state. A growing body of evidence suggests that, under appropriately controlled conditions, acute psilocybin exposure can promote long-lasting positive effects on mood and psychological well-being, offering a promising new treatment method for affective disorders (Carhart-Harris et al., 2016; Carhart-Harris & Goodwin, 2017; Davis et al., 2020; Vollenweider & Preller, 2020).

Induction of the psychedelic effect by psilocybin depends on the ability of its metabolite, psilocin (4-hydroxy-N,N-dimethyltryptamine), to act as a partial agonist of 5-HT_2A_ receptors (Vollenweider et al., 1998; Halberstadt, 2015; Madsen et al., 2019). The 5-HT_2A_ receptor is highly expressed on the dendrites of cortical layer V pyramidal neurones, particularly in the prefrontal cortex, but is also found on inhibitory interneurones and on presynaptic thalamocortical afferents (Santana et al., 2004; Saulin et al., 2012; Celada et al., 2013; Barre et al., 2016). Broadly, at the local network level, 5-HT_2A_ receptor agonists are observed to modulate glutamate transmission, disrupt typical modes of activity, and facilitate recurrent excitation (Beïque et al., 2007; Wood et al., 2012; Marek, 2018). At the systems level, human neuroimaging studies with psychedelics identify widespread disruptions to thalamocortical (Müller et al., 2017; Preller et al., 2018; Riga et al., 2018) and cortico-cortical connectivity, leading to alterations of classical functional connectivity networks and changes in neuronal dynamic properties (Kometer et al., 2013; Muthukumaraswamy et al., 2013; Carhart-Harris et al., 2014; Carhart-Harris, 2018; Mason et al., 2020; Preller et al., 2020). However, which observed neuronal effects are specific to, and characteristic of, the psychedelic state, remains to be further dissected (Roseman et al., 2014; Müller et al., 2020).

The activation of 5-HT_2A_ receptors and induction of a psychedelic state is widely suggested to promote neuroplasticity, which is theorised to be important for psilocybin’s therapeutic efficacy (Vollenweider & Kometer, 2010). While evidence exists *in vitro* and in rodents that psychedelics can induce structural and functional synaptic plasticity (Berthoux et al., 2019; Ly et al., 2018), facilitate learning and memory (Catlow et al., 2013; Zhang et al., 2013; Rambousek et al., 2018) and exert long-lasting behavioural effects (Cameron et al., 2018; Hibicke et al., 2020), the specific underlying neurophysiology remains unclear, especially with regard to the treatment of psychiatric disorders in humans.

Neuronal plasticity processes, at both structural and functional levels, whether adaptive or homeostatic, are shaped by ongoing neuronal activity. Importantly, the brain-wide changes in neuronal dynamics which occur in association with the sleep-wake cycle are strongly implicated in the regulation of plasticity of cortical function; for example, cellular maintenance, synaptic scaling, firing rate homeostasis and systems-level memory consolidation are enabled by the sleep state (Vyazovskiy & Harris, 2013; Rasch & Born, 2013; Tononi & Cirelli, 2014; Watson et al., 2016; Levenstein et al., 2017; Pacheco et al., 2020). Alongside the circadian rhythm, the occurrence of sleep is itself regulated by a homeostatic principle; prolonged wakefulness is compensated for by increased sleep intensity, enabling an approximately constant sleep quantity to be obtained each day (Borbély et al., 2016). The brain’s level of homeostatic sleep need is widely recognised to be reflected in the average levels of non-rapid eye movement (NREM) sleep slow wave activity (0.5-4 Hz) in neurophysiological field potentials (Daan et al., 1984; Achermann et al., 1993; Huber et al., 2000), such as can be observed in the electroencephalogram (EEG) or intracortical local field potential (LFP). However, slow wave amplitude and dynamics across the cortical surface reveal a heterogeneity and dependence on both behaviour and neuronal activity levels during previous wakefulness, with evidence for a bidirectional relationship between local neuronal activity and sleep-wake homeostasis (Huber et al., 2004; Rattenborg et al., 2012; Fisher et al., 2016; Thomas et al., 2020; Milinski et al., 2020).

Sleep disturbances and dysregulation are strongly associated with the development and maintenance of many common psychological disorders, including depression (Steiger & Kimura, 2010; Wulff et al., 2010; Baglioni et al., 2011; Meerlo et al., 2015). The depressed state has been theorised to be characterised by an impairment in sleep homeostasis, for example that the need for sleep increases more slowly during wakefulness and so is chronically low in depressed patients (Borbely & Wirz-Justice 1982; Wirz-Justice & Van den Hoofdakker, 1999). It is unclear whether changes in sleep architecture in depression represent simple impairments (symptoms of the disease) or instead reflect adaptive mechanisms which develop to counteract the pathophysiology of depression. The latter possibility may explain why, somewhat paradoxically, acute sleep deprivation exerts a rapid anti-depressive effect; one night of total of sleep deprivation was reported to alleviate low mood in approximately 60% of depressed patients (Wirz-Justice et al., 2005), but without additional treatments, a relapse in depressive symptoms typically occurs after subsequent sleep.

Currently very little is known about the effects of psychedelic substances on the regulation of sleep and there is a striking absence of literature exploring the possibility that the enduring beneficial effects of serotonergic psychedelics are sleep-dependent (Froese et al., 2018). There does exist evidence that psilocybin alters sleep in humans (Dudysová et al., 2020) however, animal models will be necessary to understand the underlying mechanisms. Elucidating the relationship between the actions of psychedelic serotonergic agonists and the regulation of sleep may yield insights into the core plasticity mechanisms involved in the aetiology of, and recovery from, disordered brain states such as depression.

Here, we characterise acute and enduring changes to sleep-wake-related behaviour and electrophysiology in mice following injection of psilocin, in both an undisturbed and a sleep deprived condition. We found that psilocin acutely disrupted sleep maintenance and promoted quiet wakefulness. This state was associated with altered power spectra in frontal and occipital EEG derivations and in LFPs targeted in and around the prefrontal cortex, notably including the enhancement of a 3-5 Hz rhythm and reduction in gamma band power. Despite the acute sleep disturbance, psilocin administration was not associated with long-term changes to sleep-wake architecture. After 4 hours of sleep deprivation paired with psilocin exposure, no difference was observed in slow wave activity at the EEG level, however a slower rate of slow wave activity recovery was found in the LFP of psilocin injected mice.

## Methods

### Surgical Procedures

Eight young adult male C57BL/6J mice (aged 14 - 20 weeks) were surgically implanted with electrodes for the continuous recording of electroencephalography (EEG) and electromyography (EMG), as well as with either a microwire array (n=4) or single-shank electrode (n=4) targeting the medial prefrontal cortex.

All procedures were performed under a UK Home Office Project License and conformed to the Animals (Scientific Procedures) Act 1986. Surgeries were performed under isoflurane anaesthesia (4% induction, 1 - 2% maintenance). Analgesics were administered immediately before surgery (5 mg/kg metacam and 0.1 mg/kg vetergesic, subcutaneous) and for at least three days following surgery (metacam, oral). In addition, an immunosuppressant was given both the day before surgery (0.2 mg/kg dexamethasone, intraperitoneal) and immediately before surgery (0.2 mg/kg dexamethasone, subcutaneous).

Custom-made head stages for the recording of EEG and EMG were constructed in advance of each surgery which comprised three EEG bone screws and two stainless steel EMG wires, soldered to an 8-pin surface mount connector (Pinnacle Technologies Inc., Kansas, USA). The EEG screw electrodes were inserted into holes drilled into the skull (0.7 mm drill bit, InterFocus Ltd., Cambridge, UK). One EEG screw was located above the right frontal cortex (primary motor area: anteroposterior 2 mm, mediolateral 2 mm), one above the right occipital cortex (primary visual area: anteroposterior 3.5 mm, mediolateral 2.5 mm), and one above the left cerebellum, which served as the reference signal. The two EMG wires were inserted into left and right nuchal muscle. An additional screw was located above the left occipital cortex, which served as a ground for the intracortical electrodes.

Four animals were implanted with a single-shank probe in left anterior medial cortex, aiming to span cingulate, prelimbic and infralimbic cortex (anteroposterior 1.7 mm, mediolateral 0.25 mm, depth 2 mm). The probe comprised a 5 mm shank containing 16 iridium electrode sites of 30 μm diameter, regularly spaced 50 μm apart and extending up to 800 μm from the probe’s tip (A1×16-5mm-50-703, NeuroNexus, Michigan, USA). The remaining four animals were implanted with a custom-designed polyimide-insulated tungsten microwire array (Tucker-Davis Technologies Inc., Florida, USA), spanning a larger area of left anterior medial cortex (centred anteroposterior 2.23 mm, mediolateral 0.75 mm, depth 2.2 mm, rotation 10 degrees). The array comprised 16 wire channels of 33 μm diameter arranged in 2 rows of 8, with row separation 375 μm, columnar separation 250 μm, and tip angle 45 degrees. Each wire was a custom-specified length for precise targeting of prefrontal regions (lateral row from anterior to posterior: 3.2 mm, 3.5 mm, 3.5 mm, 3.8 mm, 3.8 mm, 4 mm, 4 mm, 4 mm; medial row from anterior to posterior: 2.5 mm, 2.8 mm, 3 mm, 3.2 mm, 3.2 mm, 3.5 mm, 3.5 mm, 3.5 mm). For the single-shank probe, an additional hole was drilled to the size of the probe. For the arrays, a 1 x 2.25 mm craniotomy window was drilled into the skull. Once the array/probe was implanted, a silicone gel (KwikSil, World Precision Instruments, Florida, USA) was applied to seal the craniotomy and protect the exposed brain. Dental acrylic was used to stabilise the implanted electrodes (Super Bond, Prestige Dental, Bradford, UK) and to protect the exposed wires (Simplex Rapid, Kemdent, Swindon, UK).

### Animal Husbandry

Following surgery, mice were housed in separate individually ventilated cages and their recovery was closely monitored. The weight, spontaneous behaviour, provoked behaviour, respiration rate and grimace of each mouse was scored daily, until the mice reached baseline level for three consecutive days. Following recovery, the animals were rehoused in individual custom-made transparent plexiglass cages (20.3 × 32 × 35 cm), placed inside ventilated sound-attenuated Faraday chambers (Campden Instruments, Loughborough, UK). A camera was mounted inside each chamber and video was recorded continuously during the light period. EEG and EMG head stages were connected to the recording equipment using custom-made cables, and both LFP probe types were connected using spring-wrapped Zif-clip head stages (Tucker-Davis Technologies Inc., Florida, USA). Mice were habituated to their cables for three days before the first baseline recording began. The recording room was kept on a 12 - 12 hour light-dark cycle (lights on at 9 am), at 22 ± 1 °C and 50 ± 20 % humidity. Food and water were provided *ad libitum* throughout.

### Experimental Design

Once fully recovered from surgery, each batch of animals underwent four injection experiments, comprising all combinations of two sleep-wake conditions and two drug treatments. In one sleep-wake condition, mice were immediately returned to their home cage after injection and left undisturbed. In the second condition, a sleep deprivation protocol was enforced for four hours after injection. A within-subjects design was employed such that each mouse experienced all combinations of psilocin vs. vehicle and undisturbed vs. sleep deprivation conditions exactly once. The order of drug treatments and sleep-wake conditions was counterbalanced across all eight animals. Each injection experiment was separated by three days, following a pattern of baseline day, injection day, and recovery day.

### Preparation of Psilocin

Psilocin (4-hydroxy-N,N-dimethyltryptamine, LGC Standards) was administered by intraperitoneal injection at a dose of 2 mg/kg. Crystalline psilocin was dissolved in 50 mM tartaric acid and subsequently diluted in saline (5% glucose) up to a concentration of 0.25 mg/ml.

### Sleep Deprivation

Sleep deprivation was performed using the well-established gentle handling procedure (Fisher et al., 2016). During this period, experimenters constantly monitored both the behaviour and ongoing neurophysiological recordings of the mice. As soon as any animal showed signs of sleepiness (such as immobility, or slow waves in the EEG), novel objects were introduced to the cage (such as cardboard, colourful plastic, sponge, tin foil, wooden blocks and plastic wrap) in order to encourage wakefulness. In some cases, towards the end of the sleep deprivation period, the mice stopped responding to novel objects and were awakened instead by gentle physical stimulation (brushing with tissue paper or disturbance of their nest), although this was kept to a minimum. Sleep deprivation lasted 4 hours.

### Data Acquisition

Electrophysiological signals were acquired using a multichannel neurophysiology recording system (Tucker-Davis Technologies Inc., Florida, USA). Signals were managed and processed online using the software package Synapse (Tucker-Davis Technologies Inc., Florida, USA). All signals were amplified (PZ5 NeuroDigitizer preamplifier, Tucker-Davis Technologies Inc., Florida, USA), filtered online (0.1 – 128 Hz) and stored with a sampling rate of 305 Hz. In addition, the raw LFP signal was processed to extract extracellular multi-unit spiking. The signal was filtered (300 Hz – 1 kHz) and an amplitude threshold was manually selected for each channel to detect the occurrence of spikes. When threshold crossing occurred the time stamp and signal snippets (46 samples at 25 kHz, 0.48 ms before and 1.36 ms after threshold crossing) were stored.

Signals were read from tank formats into Matlab (using the software package *TDTMatlabSDK*) and filtered with a zero-phase 4^th^ order Butterworth filter (using Matlab functions *butter* and *filtfilt*) between 0.5 - 100 Hz for EEG/LFP signals and between 10 – 45 Hz for EMG, then resampled at 256 Hz (using the Matlab function *resample*). Spiking activity from each channel was first cleaned offline for artefacts using the Matlab spike sorting software *Wave_clus* (Quiroga et al., 2004). Although this software outputs putative sorted single units, very few separable clusters were typically found (often only one) and these were not of high quality (many inter-spike intervals below 3 ms refractory period), so unit clusters were merged and treated as multi-unit activity. However, the software was very useful for identifying waveforms which were not spike-like and were therefore discarded as electrical artefacts. Firing rate was calculated in epochs of 4 seconds separately for each channel. For some analyses, spike rates were normalised by expression as a percentage of the mean spike rate within the same channel and same vigilance state on the baseline day before psilocin injection in the relevant condition. Normalised spike rates could then be averaged across channels.

### Sleep Scoring

Vigilance states were scored manually by visual inspection at a resolution of 4 seconds using the software SleepSign (Kissei Comtec, Nagano, Japan). Wake was characterised by low amplitude irregular EEG and LFP signals alongside asynchronous high frequency multi-unit spiking. In contrast, NREM sleep was identifiable by the presence of high amplitude EEG and LFP slow waves coincident with synchronous spiking multi-unit off periods. REM sleep periods were identifiable by a reduced slow wave activity, increased theta power and readily distinguishable from waking by low EMG levels and sleep-wake context (Figure 1). In order to identify and exclude time periods with large amplitude artefactual deflections across LFP channels, a hybrid LFP signal was created comprising the maximum absolute value of any one LFP at each time point and plotted alongside EEG for manual artefact scoring.

**Figure 1.**
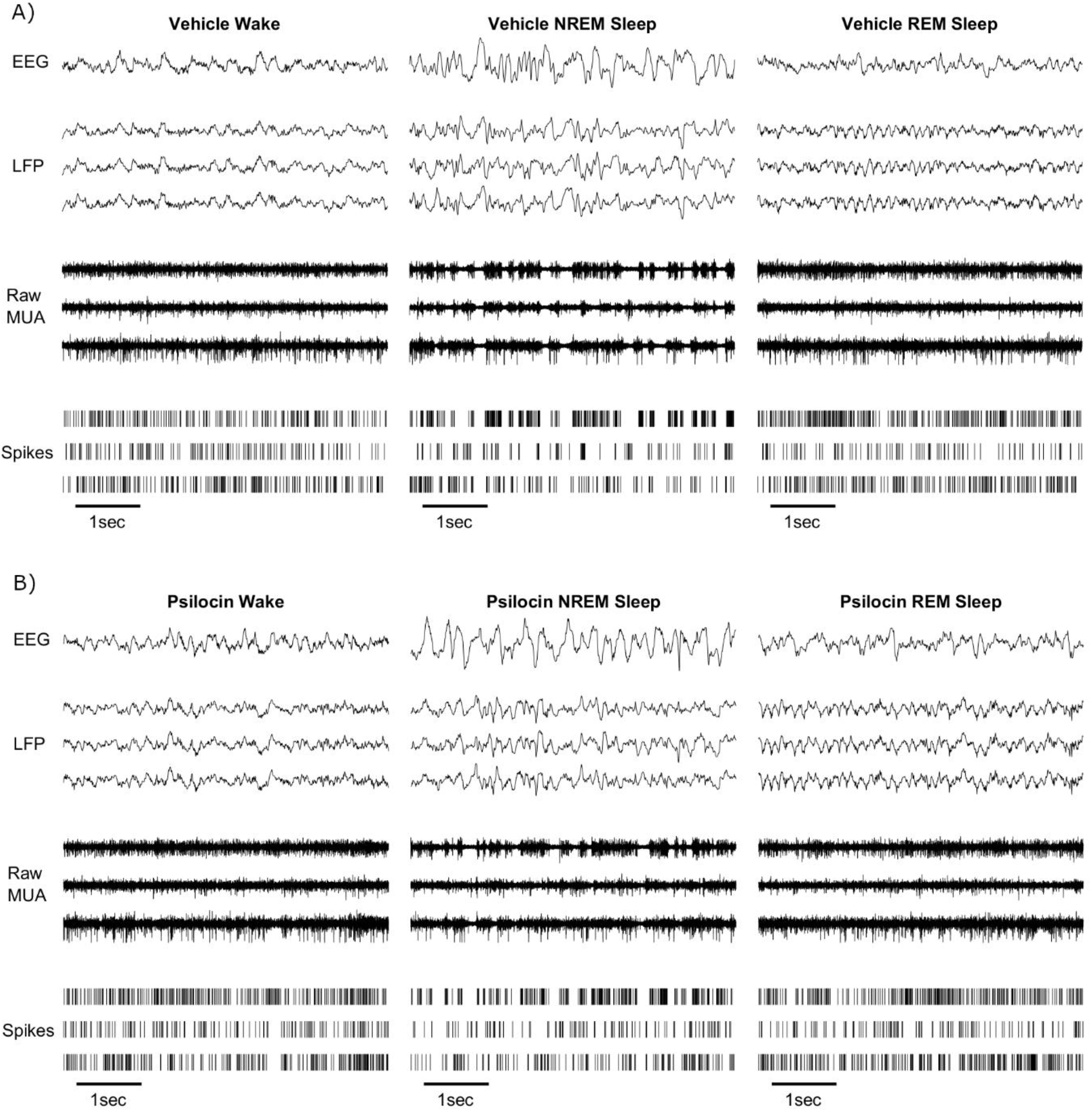
An example segment of 5 seconds duration of frontal electroencephalogram (EEG), 3 cortical local field potentials (LFP), corresponding raw signal with multi-unit activity (MUA) and detected spikes in representative segments of waking, NREM and REM sleep, soon after **A)** injection with vehicle solution, **B)** injection with psilocin.

### Histology

When all experimental recordings were complete the animals were euthanised and their brains were prepared for histological analysis for location of inserted probes. A microlesion protocol was applied immediately after death using a NanoZ stimulation device (White Matter LLC, Seattle, USA). Four channels (chosen to be as equally spaced across the area covered by the probe as possible) were sequentially stimulated with 10 μA of current for 20 seconds. Animals underwent transcardial perfusion with paraformaldehyde (PFA, 4%) and extracted brains were suspended in PFA for 24 - 48 hours before being stored in PBS (with sodium azide). Brains were sectioned into 50 μm coronal slices using a freezing microtome and stained with DAPI, a DNA binding fluorophore. Probes were stained before insertion with DiI. The slices were imaged using an Olympus FV1000 confocal microscope and compared with an anatomical atlas (Paxinos & Franklin, 2013) to aid localisation of probes.

Histology was only fully successful in four animals (two single shank and two array implanted). Based on the analysed subset, the estimated distribution of targeted cortical regions was approximately prelimbic (39%), cingulate (31%), secondary motor (12 %), medial orbital (10%) and infralimbic (8%). Histological analysis of the remaining animals suggested that some electrodes may have reached deeper and more posterior structures including the dorsal striatum and lateral septal nucleus, however this could not be definitively confirmed. For analysis, all quality LFP signals were grouped and treated as one population.

### Data Inclusion/Exclusion

Out of the 8 animals in the study, frontal EEG recordings were successfully obtained from 7, occipital EEG from 5, LFP from 6 and multi-unit spiking activity from 7. Lost signals were due to damage to the electrode or connecting wires. All signals were obtained simultaneously from 5 animals. All individual LFP signals were manually examined and a total of 15 (out of 112) were identified for exclusion based on the presence of frequent high amplitude artifacts or unsystematic drift in signal amplitude during key analysis time windows.

### Field Potential Spectral Analysis

The spectral properties of EEG and LFP signals were analysed with a discrete fast Fourier transform (FFT) on segments of 4-second duration, applying a Hann window. Spectral power values were averaged over epochs scored as the same vigilance state within the time window of interest separately for each animal. An average power for each discrete frequency value in wake, NREM and REM sleep was calculated for each animal from the whole 24-hour baseline day before psilocin injection. For plotting spectra, values between 45 - 55 Hz were interpolated for ease of visualisation, since power in this frequency range was removed by a notch filter targeting 50 Hz line noise. When specifically analysing slow wave activity, this was obtained for each epoch by summing power over frequencies from 0.5 – 4 Hz obtained from the FFT.

### Statistics

Statistical tests were all performed using Matlab. ANOVA was performed using the Matlab function *anova2* and paired samples t-tests were conducted using the Matlab function *ttest*, after confirming that data pass a Lillie’s test for normality (function *lilliestest*) and in some cases, where indicated, tests were run after applying a log transform to improve data normality. A Wilcoxon test was performed (function *ranksum*) for highly skewed data. When testing for differences between power spectra t-tests were run at all individual discrete frequency values, applying a p < 0.05 significance threshold without correction for multiple comparisons.

## Results

In this study 8 mice were injected with either 2 mg/kg psilocin (4-hydroxy-N,N-dimethyltryptamine) or vehicle while EEG, cortical LFPs and neuronal activity were continuously monitored over a period of 11 days, comprising 4 injection experiments. The within-subject design incorporated two sleep-wake conditions (undisturbed vs. sleep deprivation) and two treatments (drug vs. vehicle).

### Sleep is acutely destabilised and fragmented after psilocin

In the first condition, the mice were left undisturbed in their home cage after injection, in order to study arousal and spontaneous sleep-wake behaviour. The injection was administered shortly after light onset, at which time the animals are typically asleep. The animals were, of course, necessarily awakened by the intraperitoneal injection procedure. The most striking effect of the psilocin was to disrupt the animals’ first attempts at initiating sleep. When affected by the drug, the mice spent a significant amount of time in their nests, adopting a posture compatible with sleep, but still apparently awake according to electrophysiological criteria. Rest in these psilocin-affected animals was frequently disturbed by small body movements, such as stretches and readjustments of posture, and often the eyes remained open even while motionless (Supplementary Video 1).

Psilocin injection did not change the essential features of electrophysiological signals in wake, NREM or REM sleep in a way that was immediately visually identifiable (Figure 1), and it was therefore possible to score sleep-wake episodes in both vehicle and psilocin-treated animals. Analysis of this period using electrophysiological criteria for sleep-wake definition suggested that the psilocin-injected animals were rapidly alternating between short wake and shallow NREM sleep episodes (Figure 2A, 2B). The average latency from the time of injection to the first 4-second epoch scored as NREM sleep, based on electrophysiological criteria, was 18.6 ± 6.1 minutes in the vehicle condition and 26.3 ± 4.5 minutes in the psilocin animals, which was not significantly different between conditions (p = 0.30, n = 8, paired t-test, Figure 2C). However, the mean latency to the first NREM sleep episode, defined as continuous NREM sleep of at least 1-minute duration, was 25.7 ± 5.4 minutes in vehicle animals compared to 43.4 ± 3.7 minutes in animals injected with psilocin, which was a significant difference (p = 0.015, n = 8, paired t-test, Figure 2D). Similarly, the latency to the initiation of any REM sleep was also increased by psilocin, from an average of 44.5 ± 5.1 minutes in vehicle condition to 74.6 ± 6.1 minutes in the psilocin condition (p = 0.013, n = 8, paired t-test, Figure 2E).

**Figure 2.**
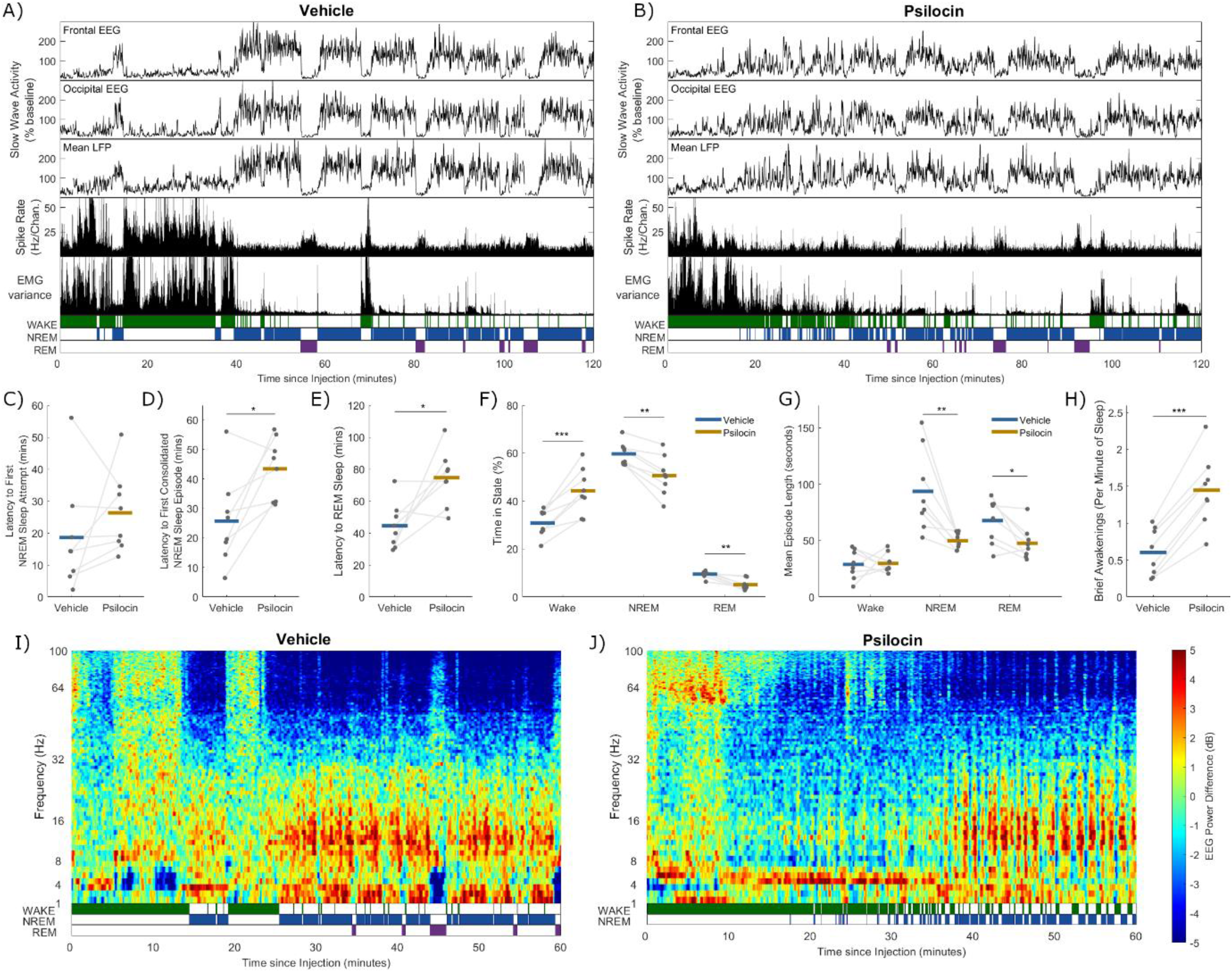
A representative example of slow wave activity (0.5 - 4 Hz power) derived from frontal electroencephalogram (EEG), occipital EEG, and mean local field potential (LFP), alongside the total recorded spike firing rate (spikes per second per channel), variance of the electromyogram (EMG) and scored vigilance states, all with a resolution of 4 seconds over a period of 2 hours after **A)** injection with vehicle, and **B)** injection with psilocin, in the undisturbed condition. The latency in minutes from injection **C)** until the first 4-second epoch scored as NREM sleep, **D)** until the first continuous NREM sleep episode at least 1-minute duration, and **E)** until the first 4-second epoch scored as REM sleep. **F)** The percentage of the three-hour period after injection which was scored as wake, NREM or REM sleep, and **G)** the mean length in seconds of wake, NREM and REM sleep episodes in this time. **H)** The number of brief awakenings (wake episodes < 20 seconds occurring within NREM episodes at least 1-minute duration) per minute of NREM sleep during the first hour after injection. In **C-H)** grey dots correspond to individual animals, with grey lines linking values from the same animal, coloured lines indicate the group mean for vehicle (blue) and psilocin (yellow) conditions and asterisks denote statistical significance with a paired t-test; *: p < 0.05, **: p < 0.01, ***: p < 0.001. Time-frequency plots characterising changes in frontal EEG spectral power (in decibels relative to power averaged over the baseline day) over the first hour after injection with **I)** vehicle and **J)** psilocin, from one representative example (different animal to **A** & **B**).

Alterations to sleep-wake activity in the psilocin-treated animals were observed to be greatest during the first hour after injection but could last up to approximately 3 hours, and so this time window was analysed further. Over the 3 hours following injection, psilocin increased the average proportion of time spent awake (Vehicle: 30.8 ± 2.0%, Psilocin: 44.3 ± 3.4%, p = 6.8 × 10^−4^, n = 8, paired t-test), and correspondingly significantly decreased the time spent in NREM (Vehicle: 59.7 ± 1.7%, Psilocin: 50.6 ± 2.9%, p = 0.0011, n = 8, paired t-test), and REM sleep (Vehicle: 9.4 ± 0.5%, Psilocin: 5.1 ± 0.8%, p = 0.0012, n = 8, paired t-test, Figure 2F). However, during this 3-hour period after injection, the mean duration of continuous wake episodes was unchanged (Vehicle: 28.8 ± 4.5 secs, Psilocin: 29.8 ± 3.1 secs, p = 0.89, paired t-test), whereas episode duration was significantly reduced for both NREM (Vehicle: 93.6 ± 12.9 seconds, Psilocin: 50.0 ± 2.6 seconds, p = 0.0070 paired t-test) and REM sleep (Vehicle: 67.7 ± 7.0 secs, Psilocin: 47.6 ± 5.1 seconds, p = 0.019, Figure 2G). This form of sleep disruption resembles an increased propensity for brief awakenings, usually defined in mice as periods of wakefulness lasting ≤ 20 seconds occurring during NREM sleep and typically accompanied by small body movements. During the first hour following injection, the frequency of brief awakenings per minute of NREM sleep was increased by psilocin (Vehicle: 0.60 ± 0.32; Psilocin: 1.4 ± 0.48; p = 2.0 × 10^−5^, n = 8, paired t-test, Figure 2H). These results suggest that the increased wakefulness produced by psilocin is due to an increased drive to awaken from sleep, corresponding to an impairment of sleep maintenance rather than an enhanced stability of wakefulness.

This period of rapidly alternating wake and NREM sleep is further illustrated in Figures 2I and 2J, showing a time-frequency plot for the frontal EEG spectral power (relative to the baseline day), from one representative example animal after both vehicle and psilocin administration. In the vehicle condition, clear vigilance state boundaries are visible in the spectrogram (Figure 2I), including wake periods with heterogenous spectral composition, NREM sleep with increased low frequency (< 30 Hz) and decreased high frequency power (> 30Hz) and REM sleep characterised by reduced low frequency (< 6 Hz) and elevated upper theta (7 – 10 Hz) power. In contrast, in the psilocin condition, the wake to sleep transition was less distinct (Figure 2J). In this example, psilocin injection is followed by approximately 10 minutes of active wakefulness characterised by elevated theta (5 – 9 Hz) and upper gamma (> 50 Hz) power. Subsequently, a quiet wake period occurs containing frequent NREM sleep attempts but dominated by wakefulness, generally characterised by reduced low frequency power (< 30 Hz) and elevated power in a narrow band around approximately 4 Hz. Approximately 35 minutes post-injection, consolidated NREM sleep becomes more distinct, indicated by increased low frequency (< 30 Hz) and decreased high frequency power (> 30Hz), although frequent brief awakenings persist. Note no REM sleep occurred in this example.

### Long-term sleep-wake architecture is unaffected by psilocin

The hour-by-hour distribution of wake, NREM and REM sleep averaged over animals for 24 hours after injection is shown in Figures 3A-C. A clear light-dark cycle is evident, in which animals are awake more throughout the dark phase, particularly during its first half (beginning between 11 and 12 hours after injection). Overall, there is no striking change in sleep-wake architecture due to psilocin on this time scale, except for the increase in wake and suppression of sleep, particularly REM sleep, in the first few hours. Importantly, there is no specific time point following the acute disruption of sleep at which a rebound in NREM or REM sleep is evident. To visualise the restoration of vigilance state homeostasis, the percentage of time since injection in each state was plotted as a function of time since injection over 24 hrs. These cumulative time courses of wake, NREM and REM sleep (Figure 3D-F) suggest that homeostasis of vigilance state quantity is restored within one day, as wake and NREM sleep quantities are no longer significantly different between drug conditions after 5 or 6 hours, and REM sleep after 12 hours.

**Figure 3.**
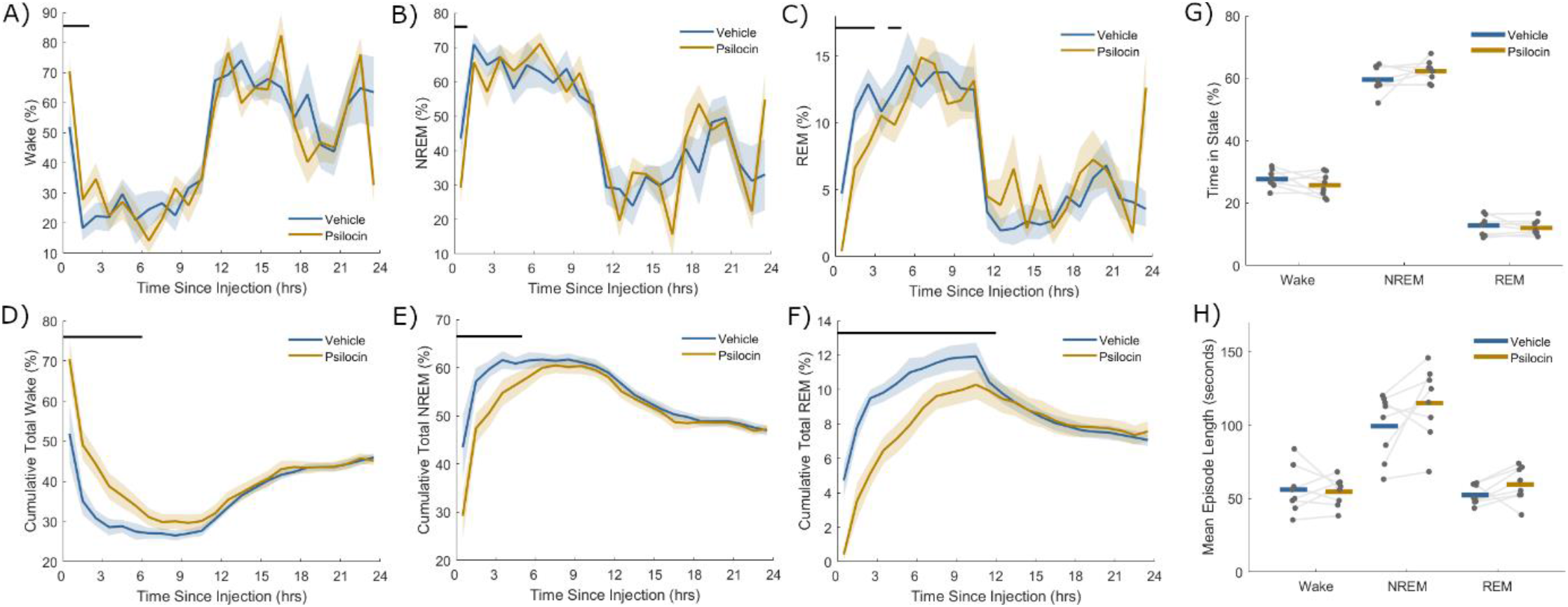
Percentage of time scored as **A)** wake, **B)** NREM sleep and **C)** REM sleep in successive non-overlapping windows of one hour up to 24 hours after injection with vehicle (blue) and psilocin (yellow). The total cumulative percentage of time scored as **D)** wake, **E)** NREM sleep and REM sleep from injection until up to 24 hours after injection with vehicle (blue) and with psilocin (yellow), as a function of time since injection. Coloured lines denote the mean and sections the standard error of the mean over all animals. Black lines indicate time points that were significantly different (p < 0.05) according to paired t-tests applied at discrete time points. **G)** The percentage time from three-hours period after injection until the end of the light period which was scored as wake, NREM or REM sleep, and **H)** the mean length in seconds of wake, NREM and REM sleep episodes in this time. Grey dots correspond to individual animals, with grey lines linking values from the same animal, coloured lines indicate the group mean for vehicle (blue) and psilocin (yellow) conditions.

Visually noticeable sleep-wake changes lasted up to three hours after injection, but from three hours after injection until the end of the light period, the total fraction of time spent in each vigilance state was not significantly different between psilocin and vehicle conditions (Wake: p = 0.25, NREM: p = 0.24, REM: p = 0.42, n = 8, paired t-test, Figure 3G). Additionally, the average duration of wake, NREM and REM sleep episodes was unchanged (Wake: p = 0.79, NREM: p = 0.13, REM: p = 0.13, n = 8, paired t-test) (Figure 3H). Similarly, the quantity of wake, NREM and REM sleep was not different in the dark period after injection (Wake: p = 0.50, NREM: p = 0.92, REM: p = 0.07, n = 4, paired t-test).

### Psilocin affects the sleep homeostatic process in a region-specific manner

The sleep-wake history of an individual is tracked by physiological processes in the brain in order to homeostatically regulate global vigilance states, such that, for example, sleep deprivation is compensated by increased subsequent sleep duration and intensity. This phenomenon is termed “Process S”, and with an underlying biological substrate that is not completely certain, measures the magnitude of the homeostatic drive to sleep, and can predict with high accuracy EEG slow wave activity through mathematical models (Daan et al., 1984; Achermann et al., 1993; Guillaumin et al., 2018; Thomas et al., 2020).

Given the pronounced acute effects of psilocin on sleep-wake states observed in the first experiment, we hypothesised that the sleep homeostatic process (Process S) would also be affected. To explore this and address the confound of psilocin’s acute direct effects on arousal, in the second experimental condition, mice were injected as before at light onset with either 2 mg/kg psilocin or vehicle, and immediately kept awake for 4 hours by engaging the animals with presentation of novel objects. We aimed to determine whether sleep quantity or slow wave activity levels would differ in subsequent recovery sleep between drug and vehicle conditions.

Overall, the electrophysiological signals during recovery sleep were similar between vehicle and psilocin conditions and the expected increased slow wave activity indicating elevated Process S was consistently observed during NREM sleep after sleep deprivation (Figure 4A, 4B). After 4 hours of sleep deprivation the median latency to the initiation of NREM sleep was not significantly different between psilocin and vehicle groups (Vehicle: 2.1 min, Psilocin: 2.2 min, p = 0.84, n = 8, Wilcoxon signed rank test, Figure 4C). There was an effect of time, but not of drug condition or interaction, on hourly sleep quantities throughout the remainder of the light period, for both NREM (Drug: F_(1,210)_ = 0.28, p = 0.60; Time: F_(14,210)_ = 6.11, p < 0.001; Interaction: F_(14,210)_ = 0.89, p = 0.57; two-way ANOVA) and REM sleep (Drug: F_(1,210)_ = 0.15, p = 0.70; Time: F_(14,210)_ = 2.16, p =; Interaction: F_(14,210)_ = 0.6, p = 0.86; two-way ANOVA).

**Figure 4.**
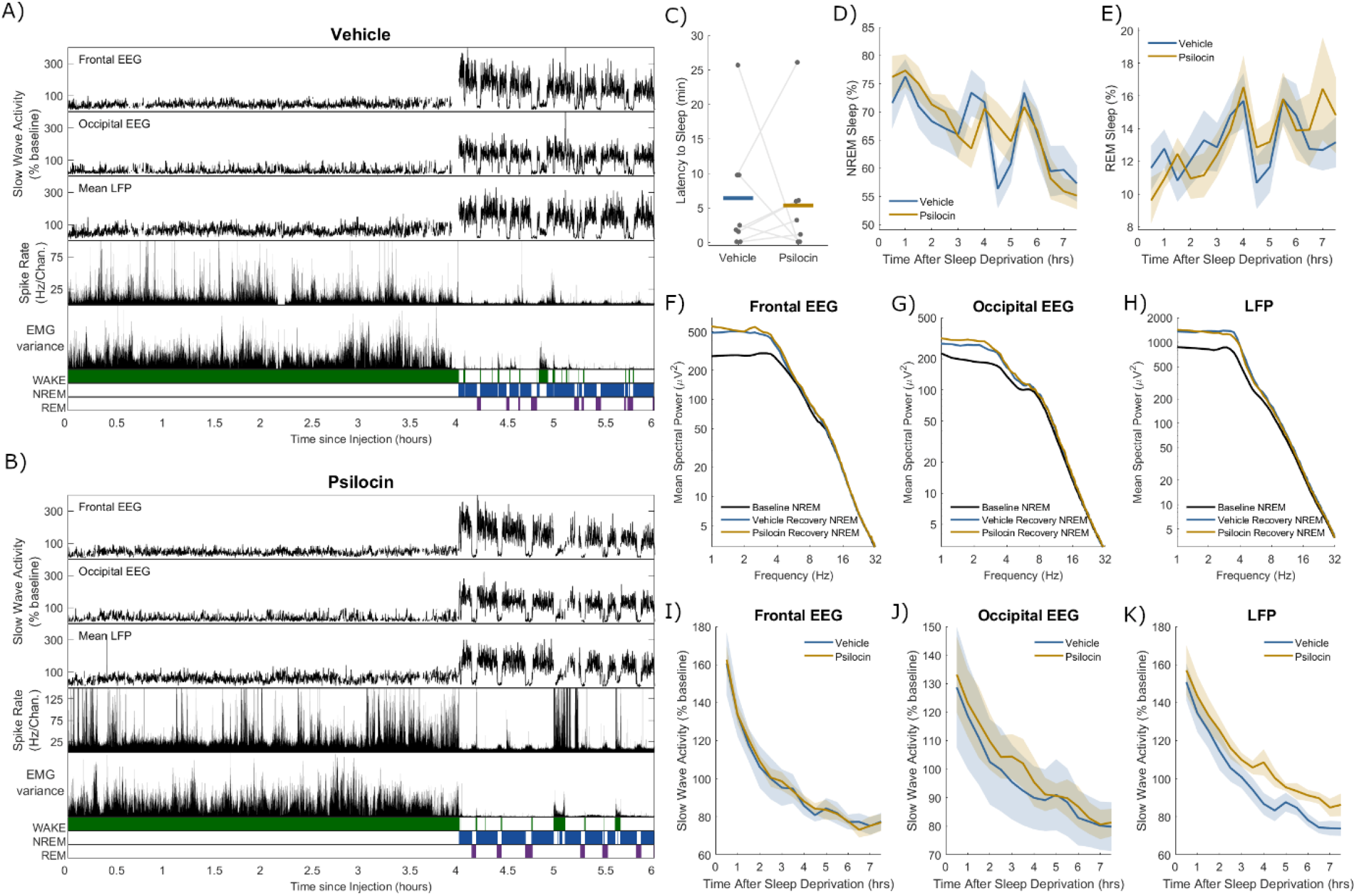
A representative example of slow wave activity (0.5 - 4 Hz power) derived from frontal electroencephalogram (EEG), occipital EEG, and mean local field potential (LFP), alongside the total recorded spike firing rate (spikes per second per channel), variance of the electromyogram (EMG) and scored vigilance states, all with a resolution of 4 seconds over a period of 6 hours comprising 4 hours of sleep deprivation and 2 hours of recovery sleep, including **A)** injection with vehicle, and **B)** injection with psilocin. **C)** Latency from the end of sleep deprivation to the first episode of NREM sleep at least 1-minute duration. Grey dots correspond to individual animals, with grey lines linking values from the same animal, coloured lines indicate the group mean for vehicle (blue) and psilocin (yellow) conditions. The percentage of time scored as **D)** NREM sleep and **E)** REM sleep from the end of sleep deprivation until the end of the light period. Coloured lines denote the mean and sections the standard error of the mean over all animals. The mean power spectra of **F)** frontal EEG, **G)** occipital EEG and **H)** mean LFP in NREM sleep in the first period of sleep after the end of sleep deprivation, compared with mean spectra over all NREM sleep the baseline day. The time series of slow wave activity (0.5 – 4 Hz power, relative to that on the baseline day) derived from **I)** frontal EEG, **J)** occipital EEG and **K)** mean LFP, from the end of sleep deprivation until the end of the light period. Individual time points correspond to averages of slow wave activity in overlapping 1-hour windows centred every 30 minutes.

While this result suggests that Process S was unaffected by psilocin administration, changes may still be visible at the level of localised cortical activity. Power spectra were calculated by averaging over NREM sleep in the first recovery sleep episode (end of sleep deprivation to first wake episode at least 5 minutes duration, Vehicle: 1.47 ± 0.7 hours, Psilocin: 1.24 ± 0.4 hours). The expected elevations in slow wave activity relative to baseline were seen in frontal EEG, occipital EEG and mean LFP, but no significant differences were observed between psilocin and vehicle conditions (Figure 4F, 4G, 4H). Furthermore, the time course of average slow wave activity in NREM sleep over the remainder of the light period after sleep deprivation further shows no effect of psilocin, only of time, in both frontal EEG (Figure 4I; Drug: F_(1,120)_ = 0.09, p = 0.76; Time: F_(14,120)_ = 27.3, p < 0.001; Interaction: F_(14,120)_ = 0.05, p = 1; two-way ANOVA) and occipital EEG (Figure 4J; Drug: F_(1,90)_ = 0.97, p = 0.33; Time: F_(14,90)_ = 3.7, p < 0.001; Interaction: F_(14,90)_ = 0.04, p = 1; two-way ANOVA). However, a significant effect for psilocin was found in the time course of mean LFP slow wave activity (Drug: F_(1,150)_ = 23.0, p < 0.001; Time: F_(14,150)_ = 24.0, p < 0.001; Interaction: F_(14,150)_ = 0.19, p = 0.99; two-way ANOVA), which exhibited a reduced rate of decrease in the psilocin condition (Figure 4K). To explore whether this result might be specific for the prefrontal cortex, it was repeated including only animals with confirmed electrode placements in prefrontal (prelimbic and infralimbic) cortex, finding the same significant effect of psilocin and a decreased decay rate of SWA during recovery sleep after sleep deprivation (Drug: F_(1,90)_ = 16.7, p < 0.001; Time: F_(14,90)_ = 17.6, p < 0.001; Interaction: F_(14,90)_ = 0.14, p = 0.99; two-way ANOVA). This result implies that the Process S recovered more slowly after psilocin, but only on a local level in prefrontal and adjacent cortex, not at the global level as measured with EEG.

### Electrophysiological characteristics of the psilocin-induced state

The effects of psilocin on EEG and LFP spectra in different states of vigilance was then explored. Wake was analysed in both the undisturbed condition (from injection until the first NREM sleep attempt, Vehicle: 18.6 ± 17.2 minutes, Psilocin: 26.3 ± 12.7 minutes) and sleep deprivation condition (first 30 minutes after injection). Both NREM and REM sleep were analysed in the undisturbed condition from the first episode of NREM sleep after injection of at least 1-minute duration, to the next wake episode at least 5 minutes duration (NREM Vehicle: 66.9 ± 22.2 minutes; NREM Psilocin: 52.2 ± 20.8 minutes; REM Vehicle: 10.1 ± 5.5 minutes, REM Psilocin: 7.3 ± 4.8 minutes). In both wake conditions, baseline spectra from frontal EEG and LFP were characterised by a peak around 4 Hz (Figure 5A, 5B, 7A, 7B). Enhancement of power around 3-5 Hz by psilocin was evidenced in the waking frontal EEG, and in the undisturbed condition in the LFP (Figure 5A, 5B, 7B). Notably this peak was reduced in the sleep deprivation condition in the LFP and with vehicle in frontal EEG (Figure 5B, 7B). Low frequency power increases in occipital EEG were broader and at a high frequency with psilocin (Figure 6A, 6B), reflecting widening of the theta (5-8 Hz) peak present in baseline spectra. These changes are likely linked to behaviour, for example the 3-5 Hz rhythm may be associated with quiet wakefulness (Contreras et al., 2021), and indeed a negative correlation was found between 3-5 Hz frontal EEG power and EMG variance per 4-second epoch as a measure of motor activity during the 3-hour period after psilocin injection (Spearman’s R = −0.47 ± 0.12, n = 5). High frequency power in the gamma range (> 30 Hz) was generally decreased by psilocin in both EEG derivations and to a greater degree in the undisturbed condition (Figure 5A, 5B, 6A, 6B). This effect was weaker in the LFP and accompanied by an increase in high frequency power (> 60 Hz) particularly during the sleep deprivation condition (Figure 7B).

**Figure 5.**
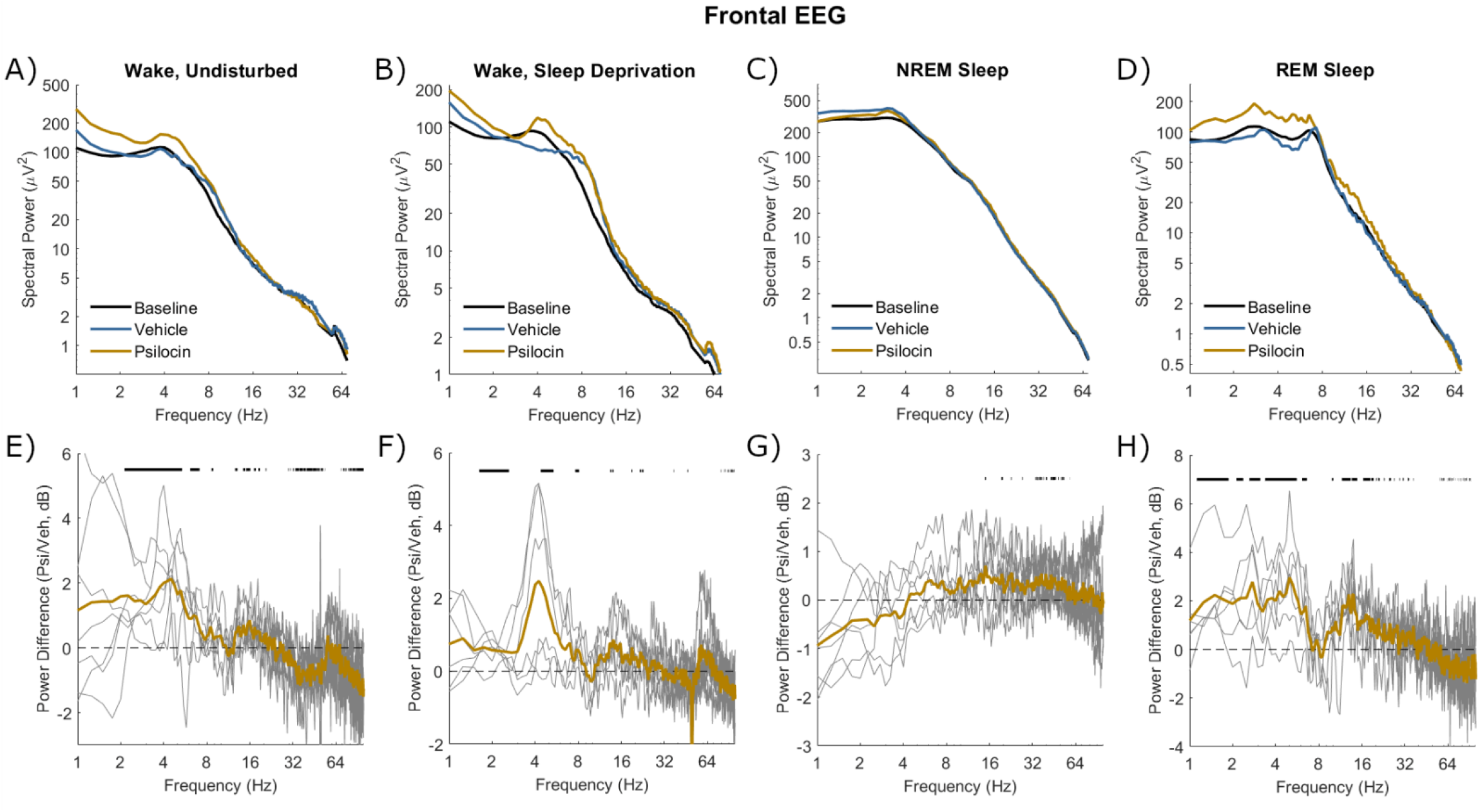
The mean power spectra of frontal EEG in four different vigilance state conditions following injection with vehicle (blue) and psilocin (yellow). **A)**‘Wake, undisturbed’ corresponds to the first experiment, all wake epochs from injection until the first NREM episode at least 1-minute duration. **B)**‘Wake, sleep deprivation’ corresponds to the first 30 minutes of sleep deprivation in the second experiment. **C)**‘NREM sleep’ and **D)**‘REM sleep’ correspond to the first experiment, all NREM/REM sleep epochs in the sleep period after injection, defined from the start of the first NREM sleep episode at least 1 minute duration until the next wake episode at least 5 minutes duration. Spectra averaged across the same vigilance state in the baseline day are shown for comparison (black). **E-H)** Below each plot illustrates the spectral power difference as a function of frequency in decibels between vehicle and psilocin conditions (positive is greater after psilocin). Grey lines correspond to individual animals and coloured lines to the mean. Black lines indicate discrete frequencies (at 0.25 Hz resolution) that were significantly different (p < 0.05) according to paired t-tests.

**Figure 6.**
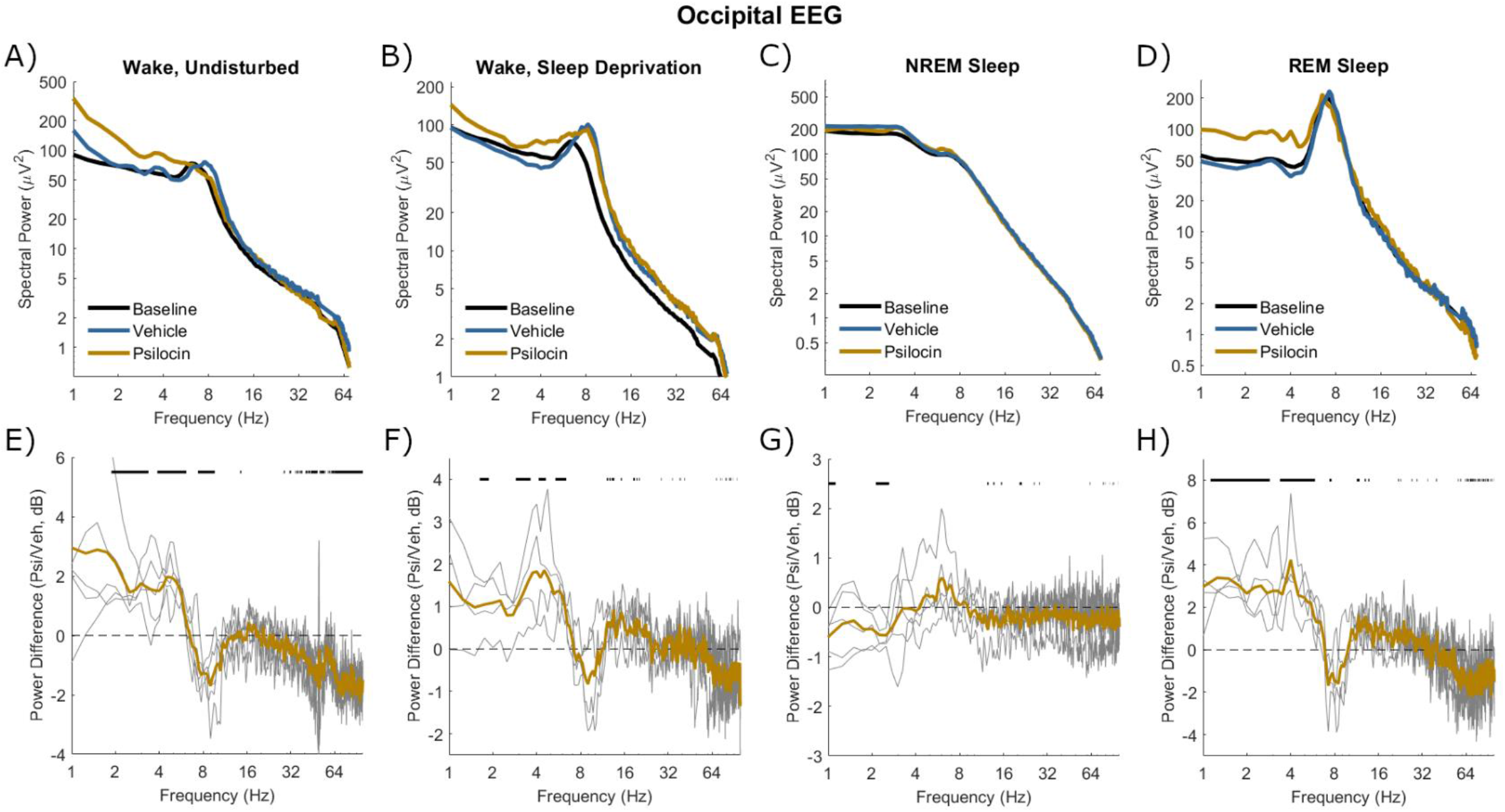
The mean power spectra of occipital EEG in four different vigilance state conditions following injection with vehicle (blue) and psilocin (yellow). **A)**‘Wake, undisturbed’ corresponds to the first experiment, all wake epochs from injection until the first NREM episode at least 1-minute duration. **B)**‘Wake, sleep deprivation’ corresponds to the first 30 minutes of sleep deprivation in the second experiment. **C)**‘NREM sleep’ and **D)**‘REM sleep’ correspond to the first experiment, all NREM/REM sleep epochs in the sleep period after injection, defined from the start of the first NREM sleep episode at least 1 minute duration until the next wake episode at least 5 minutes duration. Spectra averaged across the same vigilance state in the baseline day are shown for comparison (black). **E-H)** Below each plot illustrates the spectral power difference as a function of frequency in decibels between vehicle and psilocin conditions (positive is greater after psilocin). Grey lines correspond to individual animals and coloured lines to the mean. Black lines indicate discrete frequencies (at 0.25 Hz resolution) that were significantly different (p < 0.05) according to paired t-tests.

**Figure 7.**
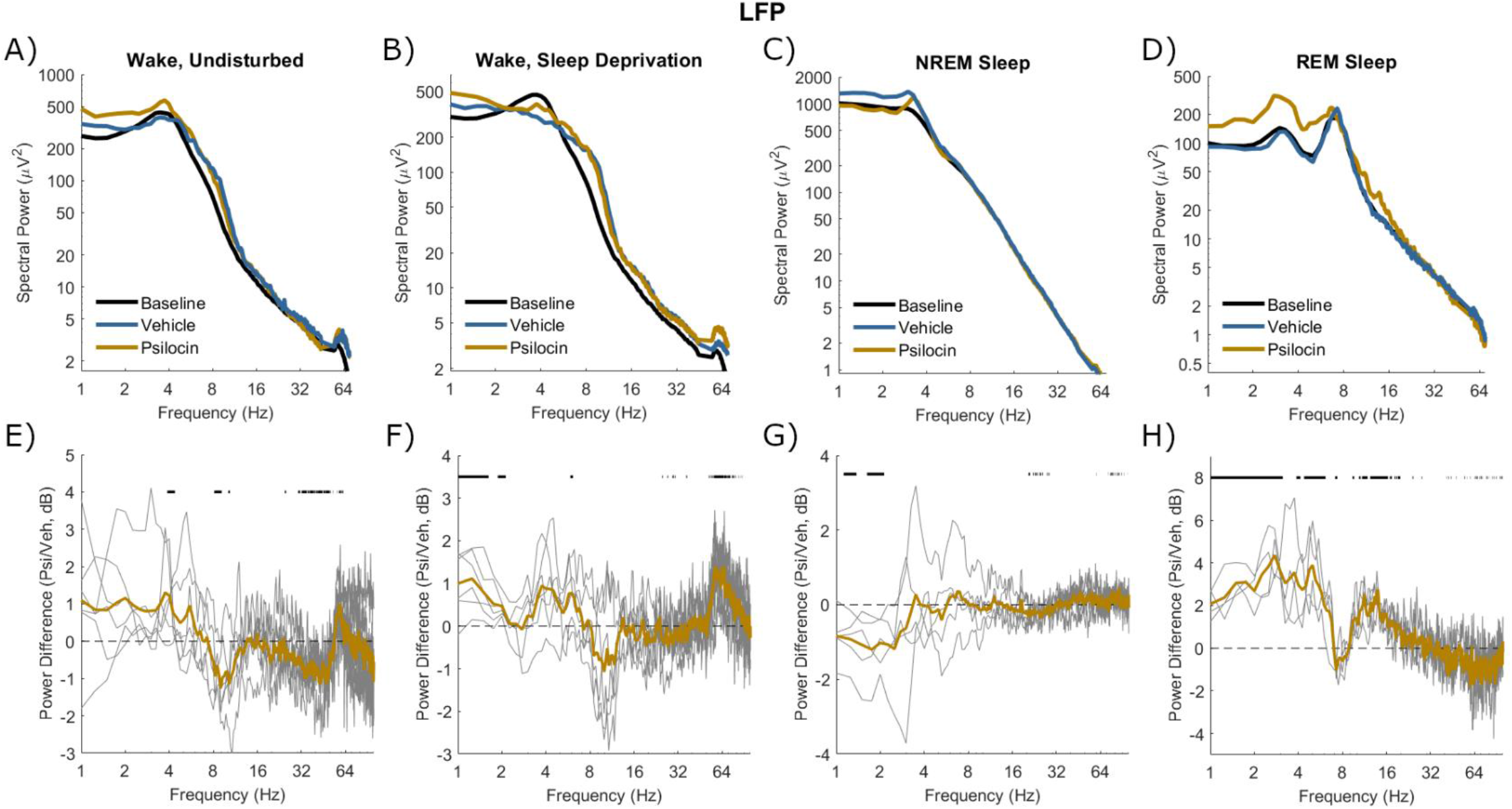
The mean power spectra of mean LFP in four different vigilance state conditions following injection with vehicle (blue) and psilocin (yellow). **A)**‘Wake, undisturbed’ corresponds to the first experiment, all wake epochs from injection until the first NREM episode at least 1-minute duration. **B)**‘Wake, sleep deprivation’ corresponds to the first 30 minutes of sleep deprivation in the second experiment. **C)**‘NREM sleep’ and **D)**‘REM sleep’ correspond to the first experiment, all NREM/REM sleep epochs in the sleep period after injection, defined from the start of the first NREM sleep episode at least 1 minute duration until the next wake episode at least 5 minutes duration. Spectra averaged across the same vigilance state in the baseline day are shown for comparison (black). **E-H)** Below each plot illustrates the spectral power difference as a function of frequency in decibels between vehicle and psilocin conditions (positive is greater after psilocin). Grey lines correspond to individual animals and coloured lines to the mean. Black lines indicate discrete frequencies (at 0.25 Hz resolution) that were significantly different (p < 0.05) according to paired t-tests.

In NREM sleep, no well-defined band-specific differences were identified between vehicle and psilocin conditions. A trend existed in all EEG and LFP signals for decreased low frequencies (< 4 Hz), perhaps reflecting that sleep was less intense (Figure 5C, 6C, 7C). During REM sleep after psilocin, high frequencies (> 30 Hz) tended to be reduced, whereas low frequencies (< 8 Hz and 10-20 Hz) were mostly increased (Figure 5D, 6D, 7D). These differences might be interpreted as a bleeding of NREM-like activities (delta waves, spindles, reduced gamma) into REM sleep.

## Discussion

The aim of this work was to explore possible effects of psilocin, a classical psychedelic and agonist of the 5-HT_2A_ receptor, on sleep-wake regulation and associated cortical activity in mice. Psilocin was administered to mice in both an undisturbed condition in which voluntary sleep occurs and a condition of enforced prolonged wakefulness. Compared to a vehicle control injection, psilocin was observed to acutely disrupt sleep, suppressing the maintenance of both NREM and REM sleep, resulting in a pattern of fragmented sleep attempts and frequent brief awakenings which lasted up to 3 hours. No enduring effects of psilocin were observed on sleep-wake quantities or episode duration. However, while the sleep homeostatic process (Process S) was not found to be disrupted by exposure to the drug in the “global” EEG signal, there was evidence for a slower decline of slow wave activity in the LFP in recovery sleep following sleep deprivation combined with psilocin injection compared to vehicle.

### Effects of serotonergic agents and antidepressants on sleep

The role of the serotonin system in sleep-wake control is complex and somewhat controversial (Ursin 2008, Monti 2011). Serotonergic raphe neurones are more active during wakefulness than sleep and serotonin is widely included in the monoaminergic ascending arousal system thought to maintain wakefulness (Saper 2010). However, optogenetic studies in mice suggest that, while burst activity in the raphe is indeed wake promoting, tonic activity contributes to the build-up of sleep need (Oikonomou et al., 2019). An acute sleep-suppressing effect of psychedelic 5-HT_2A_ receptor agonists has been previously reported in rodents and cats (Colasanti & Khazan, 1975; Kay & Martin, 1978; Monti & Jantos, 2006). Correspondingly, 5-HT_2A_ receptor antagonists are sleep-promoting in rats (Monti & Jantos, 2006) and in mice, however 5-HT_2A_ receptor knockout mice sleep less and exhibit an attenuated homeostatic sleep rebound (Popa et al, 2005). In humans, 5-HT_2A_ receptor antagonists increase the depth and maintenance of sleep and have been explored in the treatment of insomnia (Vanover & Davis, 2010; Monti et al., 2018).

REM sleep suppression is a common effect of many different antidepressants with different pharmacological profiles, including selective serotonin reuptake inhibitors, serotonin-noradrenaline reuptake inhibitors, monoamine oxidase inhibitors and tricyclic antidepressants (McCarthy et al., 2016; Wichniak et al., 2017) and REM sleep is strongly associated with the regulation of emotional memory (Perogamvros & Schwarz, 2015). Disruption of REM sleep after psychedelic drug exposure has been previously reported in humans, albeit only in the single night after drug exposure (Barbanoj et al., 2008; Dudysová et al., 2020). Similarly, the observed suppression of REM sleep in this study lasted only on the order of hours, as REM sleep levels returned to match those of vehicle controls by the end of the light period and no evidence was found for effects manifesting in the following dark period or subsequent day. Given the limited duration of effects, it is concluded that psilocin does not necessarily induce long-term changes in sleep-wake architecture in mice, at least at the 2 mg/kg dose given here, and that it is unlikely that modulation of REM sleep quantity is a core mechanism of the psychological benefits of psychedelics. It remains possible, however, that REM sleep is affected in a more subtle way in terms of the underlying network activity.

### Significance of the 3 – 5 Hz oscillation

Analysis of frontal EEG and LFP spectra in these recordings identified a prominent 3 - 5 Hz peak amplified by psilocin. This peak was present in baseline wakefulness in both conditions, so likely corresponds an enhancement of a particular form of existing activity. Oscillatory activity in this frequency range has been previously associated with breathing during wakefulness in mice in the prefrontal cortex (Biskamp et al., 2017), as well as other areas (Jessberger et al., 2016; Chi et al., 2016) and serotonergic signalling is implicated in breathing regulation (Hilaire et al., 2010). The prefrontal respiratory rhythm emerges during wake immobility, synchronises with nasal breathing and modulates ongoing prefrontal cortical gamma activity and spike timing (Biskamp et al., 2017). This respiratory rhythm has also been linked with neocortical-hippocampal communication, plasticity, and memory (Liu et al., 2017). A recent study reported periods of EEG oscillatory activity at around 4 Hz in mice after treatment with a 5-HT_2A_ receptor agonist, finding that these coincide with behavioural inactivity, although breathing was not measured (Contreras et al., 2021). That this psilocin-associated oscillation is related to respiration remains to be directly tested, and moreover its functional significance within the context of both endogenous 5-HT_2A_ receptor activation and the psychedelic phenomenon is unclear. Dissecting the behavioural and pharmacological influences on this oscillation would require more carefully controlled experiments.

### Does psilocin affect Process S?

One of the main aims of this study was to determine the effects of psilocin on Process S. Considering the ability of psilocybin to disrupt normal neuronal dynamics and to promote widespread functional plasticity, it can be predicted that psilocin injection would lead to a net increase in cortical synaptic strengths. According to the synaptic homeostasis hypothesis, this is identical to an increase in Process S (Tononi & Cirelli, 2003). However, it has been alternatively argued that the neuroplastic influence of psychedelics might actually complement or even substitute the effects of sleep, therefore reducing Process S, since both involve a temporary relaxation of functional constraints followed by self-organised re-optimisation (Froese et al., 2018).

Since the relationship between plasticity and sleep regulation is neither straightforward nor fully understood (Frank & Heller, 2019), the absence of a change in EEG slow wave activity found here should not be interpreted as evidence that neuroplasticity does not widely occur, or that psychedelics do not affected sleep regulation. A previous study in humans reported elevated EEG slow wave activity during sleep 11 hours after ingestion of ayahuasca, a traditional psychedelic drink containing dimethyltryptamine (Barbanoj et al., 2008). However, another human study with psilocybin reported that EEG slow wave activity was suppressed in the first cycle of NREM sleep (Dudysová et al., 2020). These differences may be simply due to species or drug dose, or even the possibility that the compound remained in the system at sleep onset, exerting acute effects on arousal and the expression of the sleep slow wave itself, confounding inference into Process S per se. In this study a sleep deprivation duration was chosen which exceeded the duration of observable acute effects of the drug wake and NREM sleep. It is possible that the duration of sleep deprivation was too long, and as such a ceiling effect on slow wave activity reduced the ability to discriminate the effects on Process S between psilocin and vehicle conditions in the frontal EEG. Of course, reducing the sleep deprivation exacerbates the risk that the acute sleep-inhibiting effects of psilocin remain present, so this would be difficult to disentangle.

In this regard, the finding of a reduced recovery rate of Process S locally in the LFP targeting prefrontal cortex is of potential importance. The global Process S which manifests in slow wave activity at the EEG level has been suggested to result from the integration across the brain of many local Processes S, which in turn each reflect the recent history of local neuronal activities (Thomas et al., 2020). If the rate of Process S recovery is slowed locally in prefrontal regions, it is possible that recovery may occur more quickly elsewhere in the brain, such as more posterior cortex. The functional significance of this result is not certain and a more in-depth mapping of Process S across the cortical surface would be valuable, however, it does imply that the recovery of neuronal homeostasis after exposure to psychedelics is in some way slower in prefrontal regions. Increased slow wave activity indicates elevated neuronal synchronisation, so this result may well be linked to neuroplasticity of functional networks and is consistent with the widely formed hypothesis that the prefrontal cortex is a key cortical region affected by psilocybin.

### Future outlook

Psychedelic drugs such as psilocin provide a novel approach to study the basic science underpinning sleep regulation, offering a means to manipulate the content of wakefulness and associated brain dynamics. Psychedelic stimulation offers important advantages compared to other manipulations of waking brain activity, such as optogenetic activation of cortical neurones, owing to its relative simplicity, physiological validity, pharmacological specificity, applicability to humans, and comprehensibility in terms of the associated conscious experience. Similarly, sleep represents an overlooked aspect of physiology in the efforts to understand how psychedelic-mediated mechanisms yield psychological benefits.

It is likely that the effects of psychedelics on arousal and sleep regulation might depend on circadian time and preceding sleep-wake history. For example, it is known in humans that the 5-HT_2A_ receptor density increases after sleep deprivation (Elmenhorst et al., 2012). These factors would be easy to control for and manipulate in future animal studies. Furthermore, the time scale over which psilocybin-associated plasticity (and putative associated effects on Process S) occurs is not known. While it is often assumed that plasticity must be induced during the acute experience, this is not necessarily guaranteed to be the case. If plasticity unfolds gradually over many days, or even selectively during sleep, this will not lead to changes visible in the EEG slow wave activity.

Perhaps the greatest challenge here will be to first better understand the extent to which the physiology of psychedelic action translates between animals and humans. This is particularly important, since therapeutic benefits in humans are dependent on appropriate psychological support, such as from a trained therapist. Identifying common mechanisms of action of psychedelics in both humans and rodents would be a great advance, and will require careful use of comparator drugs, with known pharmacological profiles and subjective effects, alongside translational tools such as EEG. Although many essential questions remain, a consistent picture of psychedelic biology is gradually forming, carrying great potential to inform neuroscience in areas spanning basic neurobiology to clinical practice.

## Funding acknowledgements

BBSRC (BB/M011224/1), MRC (MR/L003635/1)

